# Standard PBMC cryopreservation selectively decreases detection of nine clinically-relevant T-cell markers

**DOI:** 10.1101/2021.05.18.443634

**Authors:** Christophe M. Capelle, Séverine Cire, Wim Ammerlaan, Maria Konstantinou, Rudi Balling, Fay Betsou, Antonio Cosma, Markus Ollert, Feng Q. Hefeng

## Abstract

Biobanking is an operational component of various epidemiological studies and clinical trials. Although peripheral blood is routinely acquired and stored in biobanks, the effects of specimen processing on cell composition and clinically-relevant functional markers of T cells still require a systematic evaluation. Here we assessed 25 relevant T-cell markers and showed that the detection of nine membrane markers, e.g., PD-1, CTLA4, KLRG1, CD25, CD122, CD127 and others reflecting exhaustion, senescence and other functions was reduced among at least one T-cell subset following standard processing, although the frequency of CD4, CD8 and regulatory T cells was unaffected. Nevertheless, a six-month-long cryopreservation did not impair the expression levels of many other membrane and all the eight tested intracellular lineage or functional T-cell markers. Our findings uncover that several clinically-relevant markers are particularly affected by processing and the interpretation of those results in clinical trials and translational research should be done with caution.

## Introduction

Biobanks substantially contribute to the implementation of different types of clinical trials and translational research in all disease areas (Baker, 2012; De Souza and Greenspan, 2013; Hewitt, 2011; Peeling et al., 2020). Each year, millions of different types of samples are being collected, stored and redistributed to perform biochemical, histopathological or genomics, proteomics and other types of “omics” analysis (Bycroft et al., 2018; Kinkorova, 2015), which is particularly important in the era of precision and personalized medicine. Among others, whole blood or isolated peripheral blood mononuclear cells (PBMCs) are routinely collected and stored for either disease-focused cohorts or vaccine trial studies (Sambor et al., 2014). The general cryopreservation effects have been long observed (Cao et al., 2003) and recently reviewed (Betsou et al., 2019). Meanwhile, more precisely no effect has been observed in the percentages of major immune subsets, such as CD4 T cells, CD8 T cells, B cells, CD14+ monocytes and CD56+ natural killer (NK) cells following cryopreservation (Anderson et al., 2019; Tompa et al., 2018). The frequency of Tregs and effector CD8 T cells has also been found stable (Gomez-Mora et al., 2020). Interestingly, the functional responses exhibited more contradictory results. For instance, proliferation responses and cytokine production in supernatants following cryopreservation of total PBMC were found to be either maintained (Carollo et al., 2012; Chen et al., 2020; Kreher et al., 2003; Reimann et al., 2000), enhanced (Anderson et al., 2019) or decreased (Axelsson et al., 2008). After cryopreservation, a reduced cytokine production was observed in CD4 T cells (Ford et al., 2017) and more recently also in antigen presenting cells (Martikainen and Roponen, 2020). Additionally, a decrease in the expression of a selected marker of interest, e.g., PD-1 in CD8 T cells was observed following cryopreservation, despite well-established cryopreservation practice in biobanking settings (Campbell et al., 2009). Several modifications in NK-cell activation markers, such as CD25, CD69 and NKP46 have been reported in frozen versus fresh samples (Gomez-Mora et al., 2020). In short, progress has already been made in assessing the effects of cryopreservation on the expression of sporadic single markers of interests in T cells. However, no work has systematically investigated the effects of cryostorage on lineage and functional T-cell markers to support evidence-based protocols for clinical trials or research studies. To this end, we analyzed up to 25 different T-cell related markers by comparing fresh and cryopreserved PBMCs from six different healthy donors, using multiple panels of state-of-the-art multi-color flow cytometry. We analyzed the markers defining T cell subsets, such as Tbet, GATA3, RORgt and FOXP3 for Th1, Th2, Th17 and Treg, respectively, as well as various functional markers in CD4 and CD8 T cell subsets, reflecting various functional attributes, including but not to limited to, activation, proliferation, exhaustion, senescence, memory/naïve status, metabolism and other effector functions.

## Results and Discussion

To assess whether the standard processing/cryopreservation conditions affect the detection of T-cell related markers, we used panels of multi-color flow cytometry to quantify and analyze up to 25 relevant T-cell lineage or functional markers in fresh, nine-week- or six-month-cryopreserved PBMCs (**Fig. 1a and Supplementary Table 1**). Similar to the results of others (Anderson et al., 2019; Tompa et al., 2018), the frequency of the three major T-cell subsets, CD4^+^FOXP3^-^ cells (Tconv), CD4^+^FOXP3^+^ regulatory T cells (Tregs) and total CD8 T cells, among living lymphocytes was unchanged before and after six-month cryopreservation (**Fig. 1b**). Unexpectedly, as early as nine weeks post cryopreservation, the frequency of cells expressing exhaustion, senescence or effector/activation markers, such as PD-1, KLRG1, CD127 and CD25, started to significantly decrease among CD4 Tconv and CD8 T cells (**Fig. S1a-e**). Following nine-week cryopreservation, both PD-1 and CD25 also started to decrease in the Tregs sub-population, which express relatively high levels of PD-1 and CD25 in fresh samples (**Fig. S1c**). To exclude the potential effects in binding affinities or fluorescence brightness/sensitivity due to the selected clone and/or fluorochrome, we chose another antibody (ab) for each of those affected markers to analyze six-month-frozen samples (for the selected antibodies (abs), refer to **Supplementary Table 1**). Importantly, to guarantee having comparable results between the different antibodies measuring the same markers, we mainly evaluated the percentage of the cells expressing the respective marker among the given T-cell subset. Notably, despite having used distinct ab clones, the frequency of cells expressing exhaustion, senescence or effector markers, such as PD-1, KLRG1, CD127 and CD25, still significantly decreased among all the three T-cell subsets in six-month cryopreserved PBMCs (**Fig. 1c-f, Fig. S2a**). These results indicate that the effect of cryopreservation is independent from ab clones and storage periods at least for the tested markers. Our data were concordant with the previously reported observation about decreased PD-1 expression among CD8 T cells following cryopreservation (Campbell et al., 2009). Although others have reported no effect of cryopreservation on CD127 expression (Tompa et al., 2018), the authors had focused on Tregs, where the expression of CD127 is known to be low or absent and therefore their report is not conflicting with our observations. The percentages of cells positive for the activation marker ICOS among CD4 Tconv, Tregs and CD8 T cells were also significantly decreased following six-month cryopreservation **(Fig. 1c, Fig. S2a)**, which was not yet the case after nine-week cryopreservation (**Fig. S1a,c,e**). For the Treg functional marker CD39, no significant effect was observed in nine-week frozen vs. fresh samples, regardless of the T-cell subsets (**Fig. S1a, c, e**). However, both the CD39 percentage and mean fluorescence intensity (MFI) were decreased after six-month cryopreservation in all the three tested T cell subsets, despite the low-level baseline expression of CD39 in CD4 Tconv and total CD8 T cells. (**Fig. S2b,c**).

**Figure 1.**
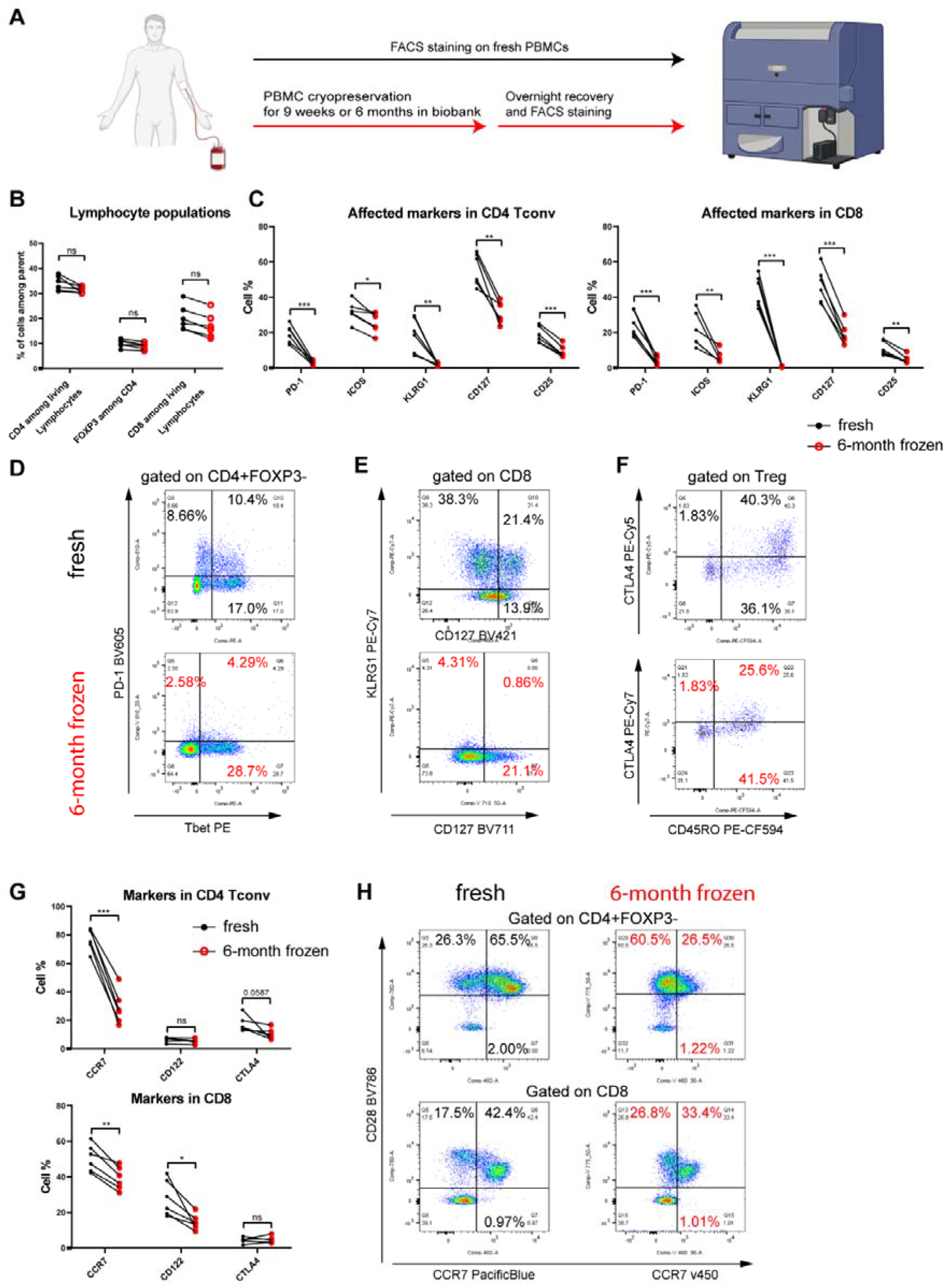
PBMC cryopreservation decreases the detection of clinically-relevant T-cell extracellular markers. **A.** Graphical representation of the experimental setup for analysing a cohort of six healthy young donors. **B.** Frequency of CD4 T cells, FOXP3+ Treg and CD8 T cells among the parent gate (living cells for CD4 and CD8, and CD4 T cells for Treg, respectively). **C.** Frequency of markers that are affected by 6-month cryopreservation in CD4 Tconv (left panel) and CD8 T cells (right, right panel). **D.** Representative flow-cytometry plots showing the co-expression of PD-1 (affected) vs. Tbet (unaffected) in fresh (black) or 6-month frozen (red) PBMCs among CD4+FOXP3-T cells (CD4 Tconv). **E.** Representative flow-cytometry plots showing the co-expression of KLRG1 (affected) vs. CD127 (affected) in fresh (black) or 6-month frozen (red) PBMCs among CD8 T cells. **F.** Representative flow-cytometry plots showing the co-expression of CTLA4 (affected) vs. CD45RO (unaffected) in fresh (black) or 6-months frozen (red) PBMCs among CD4+FOXP3+ Treg. **G.** Frequency of markers that were affected only in one or two subsets of the three T-cell subsets (CD4 Tconv, Treg and CD8 T cells). **H.** Representative flow-cytometry plots showing the co-expression of CD28 (unaffected) vs. CCR7 (affected) in fresh (black) or 6-month frozen (red) PBMCs among CD4 Tconv (above panel) and CD8 T cells (below panel). Of note, the abs used for 6-month frozen and fresh samples might be different (refer to **Supplementary Table 1**). Statistical significance was calculated by a paired Student t test. Black circles: staining on fresh PBMCs, Red circles: staining on 6-month frozen PBMCs. The value of each sample from the cohort is displayed in the figures. ns or unlabeled, not significant, *P<=0.05, **P<=0.01 and ***P<=0.001.

We also observed a certain degree of cell-type specific effects for a few markers, such as CTLA4, CD122 and CCR7. The frequency of CTLA4^+^ cells (**Fig. S2d**) was significantly decreased among Tregs and showed a similar trend among CD4 Tconv (p=0.059, **Fig. 1g**) following six-month cryopreservation. No significant effect on CTLA4 detection was observed among total CD8 T cells following either nine-week (**Fig. S1e**) or six-month cryopreservation (**Fig. 1g)**, possibly due to its low baseline expression even among fresh CD8 T cells. Unexpectedly, the frequency of CTLA4^+^ cells was significantly increased among Tregs in nine-week frozen vs fresh samples (**Fig. S1c**). The frequency of CD122^+^ cells among total CD8, but not among CD4 Tconv (**Fig. 1g)** and Tregs (**Fig. S2d**) cells, was significantly decreased following six-month cryopreservation (**Fig. 1g**). In contrast to the observation on CD122, the frequency of CCR7+ cells was significantly reduced among CD4 Tconv (**Fig. 1g, h**) and Tregs (**Fig. S2d)** following six-month cryopreservation, while that among CD8 T cells (**Fig. 1g, h**) was modestly, but still significantly decreased. After nine-week cryopreservation, the percentages of CCR7^+^ cells among CD4 Tconv and Tregs were unaffected (**Fig. S1a, c**). Nevertheless, the frequency of CCR7 expressing cells was surprisingly increased among total CD8 T cells in nine-week-old samples (**Fig. S1e**). For the nine-week cryopreserved samples, the effects on the percentages of CCR7 and CD122 positive cells again swapped, i.e., the expression of CD122 was increased among CD4 Tconv and Tregs (**Fig. S1a, c**), but remained unaffected among total CD8 T cells (**Fig. S1e**).

The freeze/thaw effects were not consistent even among all the tested cell surface markers. For instance, the frequency of other tested membrane functional markers, such as CD45RO, CD45RA, CD27, CD28, CD31, did not show any significant change among CD4 Tconv, Tregs or total CD8 T cells following six-month cryopreservation (**Fig. S3a-d**). The percentage of positive cells for the immunosenescence marker CD57 was also unaffected among both CD4 Tconv and CD8 T cells, while it showed a modest, but significant decrease among Tregs (**Fig. S3a-c**). More encouragingly, all the examined intracellular lineage or functional markers, such as FOXP3, TBET, ROGT, GATA3, EOMES, Ki-67 and pS6, were robustly detected in the corresponding positive cells, among CD4 Tconv, Tregs or total CD8 T cells (**Fig. 1b**, **Fig. S3a-c**). The frequency of natural Treg or activation marker HELIOS expressing cells was also unaffected among CD4 Tconv and total CD8 T cells, while slightly increased in Tregs, possibly due to an increased accessibility of abs to cellular nucleus proteins after cryopreservation and thawing procedures (**Fig. S3a-c**). This shows that the protein markers within cells might be better protected from cryopreservation related damage.

In summary, we identified a so far unrecognized systematic cryopreservation effect on T cells, i.e., the decrease in expression and/or detection of nine clinically-relevant markers among at least one of the three, or all three analyzed the human T-cell subsets (CD4 Tconv, Tregs and CD8 total). Those markers correspond to the critical functions of immune exhaustion, immunosenescence, activation or effector functions of T cells. Meanwhile, we did not observe a clear cryopreservation effect on the expression of at least six important cell surface markers and all eight examined intracellular markers among different subsets of T cells. We also noticed a relatively smaller number of affected markers in nine-week-versus six-month-old cryopreserved samples, indicating that the cryostorage duration might be another factor mediating the detection and/or expression levels of some specific markers. Furthermore, the dynamic effects on specific markers, for instance, the nine-week cryopreservation-induced increase in the expression of CTLA4, CCR7 and CD122, followed by six-month cryopreservation-induced decrease in their expression among specific T-cell subsets, deserve even more attention and formal validation in future clinical research endeavors. These original data provide evidence about the potential preanalytical (cryopreservation) impact on the analysis of some of the most clinically-relevant lineage and functional markers among major T cell subsets. Since cryopreservation is widely accepted and utilized among most of the completed and ongoing clinical trials worldwide, our work establishes the basis to critically assess conclusions based on immunophenotypic analyses of cryopreserved PBMCs. Our results also reveal some robust intracellular and cell surface markers. These are preferred readouts in the context of future clinical trials, since they can be more easily implemented in clinical analysis processes of various diseases involving immunological dysregulations.

## Supporting information

Three Supplementary Figures and two Supplementary Tables

## Acknowledgements

F.Q.H. was partially supported by Luxembourg National Research Fund (FNR) CORE programme grant (CORE/14/BM/8231540/GeDES), FNR AFR-RIKEN bilateral programme (TregBAR, F.Q. H. and M.O.), PRIDE programme grants (PRIDE/11012546/NEXTIMMUNE and PRIDE/10907093/CRITICS for the PhD student C.C.). The work was also partially supported through intramural funding of the host institutes LIH and LCSB through Ministry of Higher Education and Research (MESR) of Luxembourg. The funders had no role in study design, data collection and analysis, decision to publish, or preparation of the manuscript.

## Author contributions

C.C. designed and performed the experiments, performed data analysis, visualization and drafted the manuscript. S.C. and W.A. performed parts of experiments. M.K. performed parts of analyses and flow cytometry quality control. A.C., R.B., M.O., F.B., provided substantial insights and supervision into the project. F.Q.H. conceived and oversaw the whole project and wrote and revised the manuscript.

## Materials and Methods

### Blood sampling, PBMC isolation and Cryopreservation

Blood collection according to Luxembourgish ethical guidelines. Six healthy volunteers donated blood after providing informed consent. Blood was collected in a single session for all participants via venipuncture into five 10 ml vacutainer K2EDTA blood collection tubes (367525, BD) per donor. The blood collection tubes were transported at room temperature (RT) to the central processing laboratory.

Within four hours of collection, five blood collection tubes per donor were pooled and distributed over four 50 ml centrifugation tubes. Blood was diluted 1x with DPBS (14190-144 Gibco, Thermo Fisher Scientific). Diluted blood was transported in 50 ml SepMate™ tubes (86450, Stemcell Technologies) pre-loaded with Lymphoprep™ (07801, Stemcell Technologies) for PBMC isolated by density gradient centrifugation. Tubes were centrifuged at 1200 g for 15 min at RT. The interphase per tube was collected in a new 50 ml centrifuge tube and washed with DPBS using centrifugation at 500 g for 10 min at 21 °C. Cell pellets per donor were pooled in a single tube and washed with AutoMACS running buffer (130-091-221, Miltenyi, Germany) using identical centrifugation conditions. Cell pellet per donor was resuspended in 20 ml AutoMCAS running buffer. Cell concentrations were determined by Cellometer (Nexcellom, England). PBMCs per donor were diluted in in cold (2-8 °C) cryopreservation media (Cryostore CS10, Biolife Solutions) at 10 million per ml. 1 ml was transferred into a 2 ml cryovial (Greiner, Belgium). The vials were frozen using a Mr. Frosty™ freezing container, placed for up to 24 h at a −80 °C freezer, before being transported to liquid nitrogen tanks for a long-term storage. Meanwhile, 10 million of PBMCs were kept in AutoMACS running buffer at 2-8 °C and used for multiple-panel analysis of baseline flow cytometry determination (i.e., fresh samples).

### Thaw conditions, flow cytometry staining and analysis

PBMCs were either stained freshly after isolation by density gradient centrifugation or following a period of cryopreservation (nine weeks or six months for this project) in the Luxembourg local biobank (IBBL). The frozen PBMCs were first recovered by washing the cells in complete IMDM medium (supplemented with 10% FBS, PenStrep, non-essential amino acids and β-mercaptoethanol) and incubated overnight at 37 °C in the same medium. The detailed information about the components of the complete IMDM medium used for human T cells was already described in our previous work (Capelle et al., 2021; Danileviciute et al., 2019). The fresh and thawed PBMCs were stained following the same protocol as follows.

The PBMCs were first resuspended in FACS buffer (PBS + 2% FBS) containing Fc blocking antibodies (BD, 564765) and incubated for 15 min at 4 °C. After one washing step (300 g, 5 min, 4 °C) in FACS buffer, the cells were resuspended in Brilliant stain buffer (BD, 563794) containing the fluorochrome-coupled antibodies (**Supplementary Table 1**) and incubated for 30 min at 4 °C in the dark. The surface staining was followed by three washing steps in FACS buffer (300 g, 5 min, 4 °C). Next, the cells were fixed for 1 h at RT using the True-Nuclear transcription Factor Buffer Set (BioLegend, 424401). Following the fixation, the cells were centrifuged down (400 g, 5 min, RT), resuspended in 200 ul of FACS buffer and left at 4 °C over night. The next morning, the PBMCs were washed once with Permeabilization buffer (400 g, 5 min, RT) and incubated in Perm buffer containing the antibodies against the intracellular targets for 30 min at RT in the dark. Finally, the cells were washed three times in Perm buffer (400 g, 5 min, RT) and resuspended in FACS buffer for the acquisition on the BD Fortessa. The analysis of the data was performed using FlowJo.

At each time point, single-stained compensation beads (UltraComp eBeads, **Supplementary Table 2**) were acquired to generate the spillover matrix and check the quality of the antibodies. Cytometer setup was performed according to manufacture instruction using the CS&T beads (BD, 656505) to monitor optimal performance of the instrument over the time.

### Ethic statement

We complied with all the relevant ethic regulations and Informed consent was obtained from each healthy subject before the blood was drawn. The blood sampling was coordinated and performed through the Clinical and Epidemiological Investigation Centre of LIH.

### Statistics and Reproducibility

We calculated all the P values using two-tailed paired Student t test implemented in Graphpad (Prism) as specified in the corresponding Figure legends. The value of each individual sample was visualized at different time points.

## Data availability

The entire dataset of the annotated FCS files will be deposited in *Flowrepository,* which will be publicly available upon acceptance.

## Supplementary Information

Three Supplementary Figures and two Supplementary Tables are provided in the Supplementary Information.

