## Supplementary material for "Standard PBMC cryopreservation selectively decreases detection of nine clinically-relevant T-cell markers": Three Supplementary Figures and two Supplementary Tables

^4^, Integrated BioBank of Luxembourg (IBBL), 1, rue Louis Rech, L-3555, Dudelange, Luxembourg

^5^, Laboratoire national de santé (LNS), 1, rue Louis Rech, L-3555, Dudelange, Luxembourg

^6^, Quantitative Biology Unit - National Cytometry Platform, Luxembourg Institute of Health, L-4354 Esch-sur-Alzette, Luxembourg

^7^, Department of Dermatology and Allergy Center, Odense Research Center for Anaphylaxis (ORCA), University of Southern Denmark, Odense, 5000 C, Denmark

^8^, Institute of Medical Microbiology, University Hospital Essen, University of Duisburg-Essen, D-45122 Essen, Germany

Supplementary Information

#### Extended Data Figures

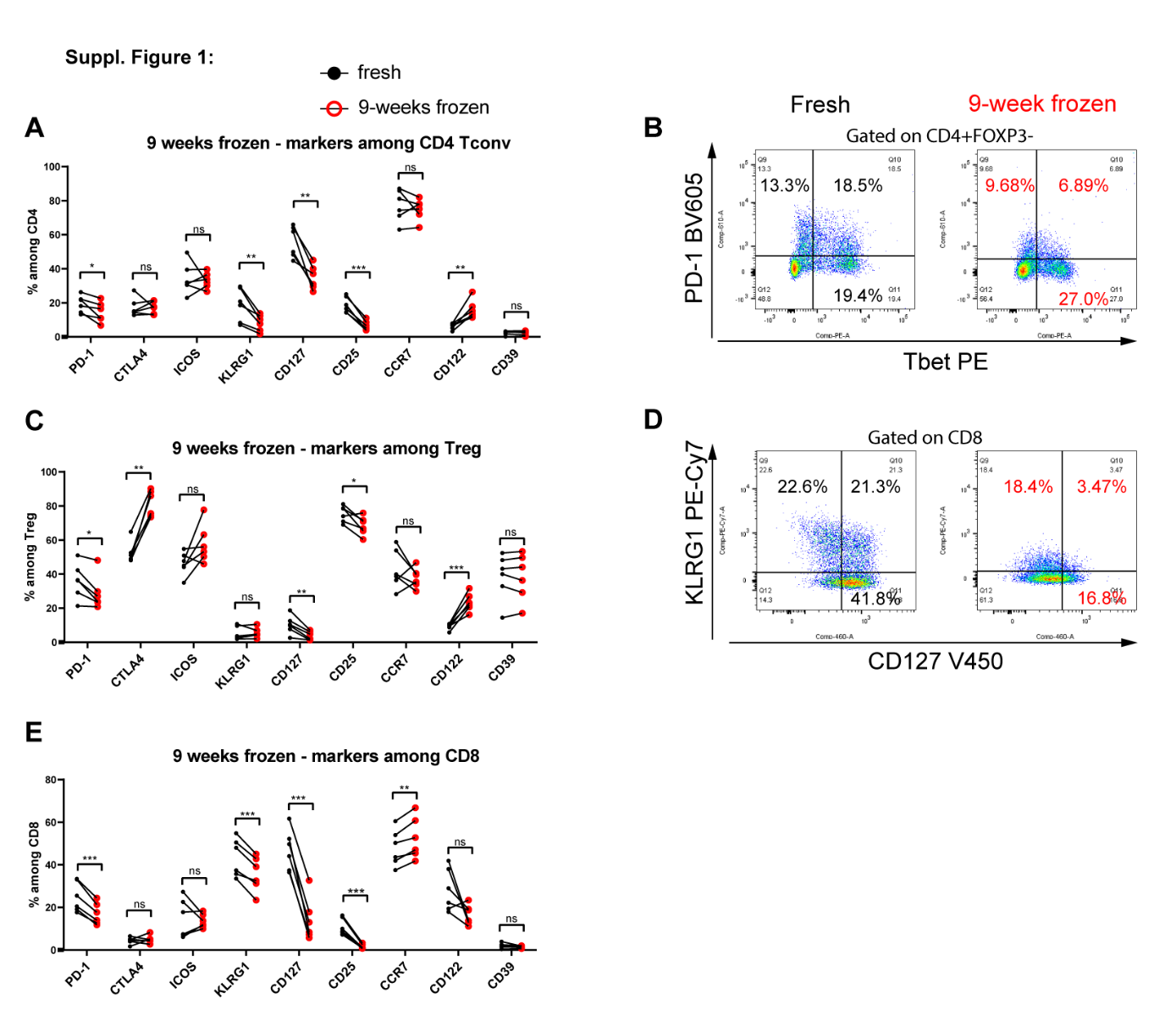

##### Extended Data Figure 1. Nine-week cryopreservation starts to decrease the detection of clinically-relevant T-cell extracellular markers.

**A.** Frequency of different markers among CD4 Tconv in fresh vs. 9-week frozen PBMCs. **B** Representative flow-cytometry plots showing the co-expression of PD-1 (affected) vs. Tbet (unaffected) in fresh (black) or 9-week frozen (red) PBMCs among CD4+FOXP3- T cells. **C.** Frequency of different markers among CD4 Treg in fresh vs. 9-week frozen PBMCs. **D.** Representative flow cytometry plots showing the co-expression of KLRG1 (affected) vs. CD127 (affected) in fresh (black) or 9-week frozen (red) PBMCs among CD8 T cells. **E.** Frequency of different markers among CD8 T cells in fresh vs. 9-week frozen PBMCs. Of note, the abs used in both 9-week frozen and fresh samples are identical. Statistical significance was calculated by a paired Student t test. Black circles: staining on fresh PBMCs, Red circles: staining on 9-week frozen PBMCs. The value of each sample is displayed in the figures. ns or unlabeled, not significant, *P<=0.05, **P<=0.01 and ***P<=0.001.

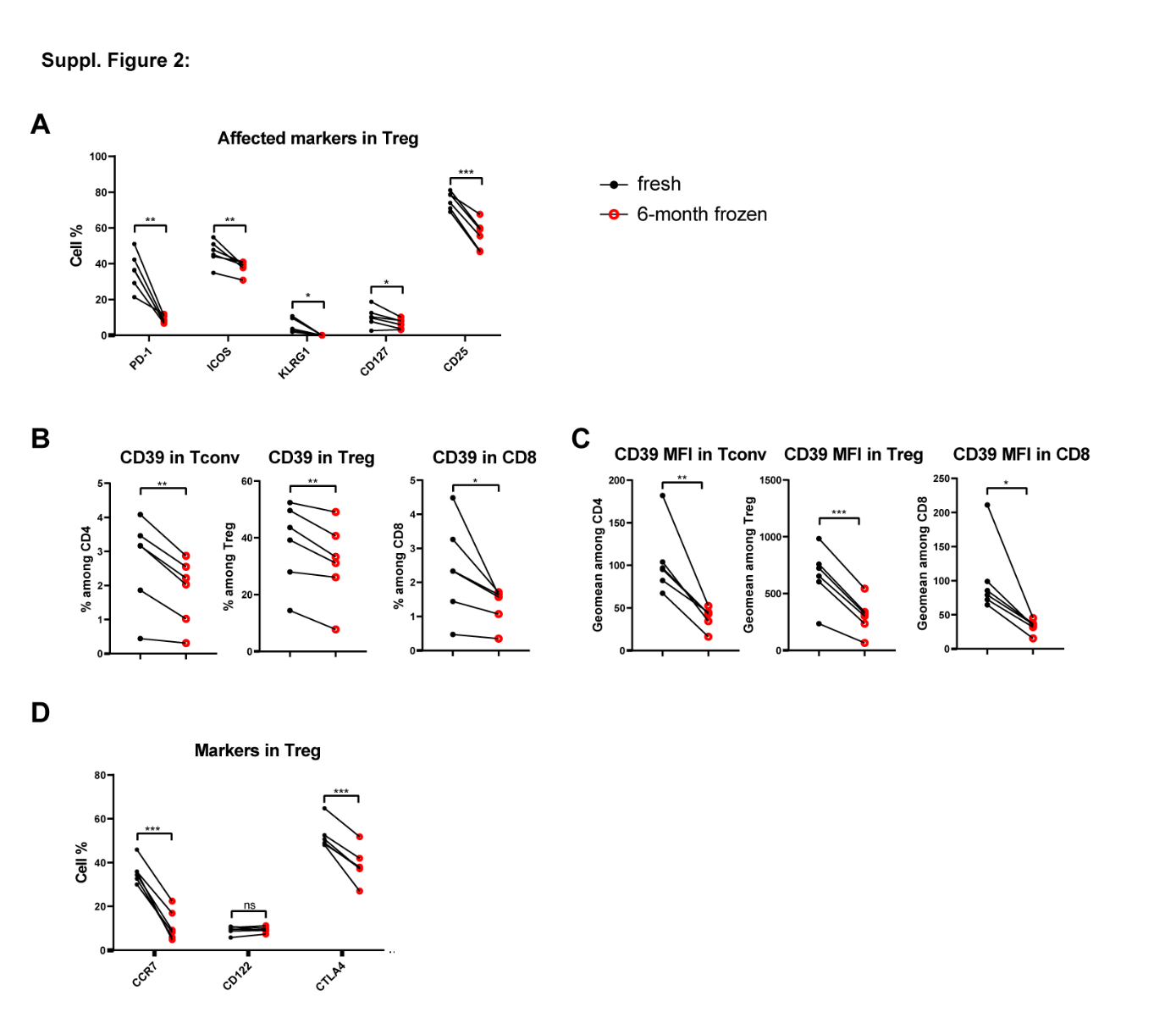

##### Extended Data Figure 2. Cryopreservation decreases the detection of other T-cell markers.

**A.** Frequency of markers that are affected by 6-month cryopreservation in CD4 Treg. **B** Frequency of CD39 in CD4 Tconv, Treg or CD8 T cells, respectively. **C.** Geomean of CD39 in CD4 Tconv, Treg or CD8 T cells, respectively. **D.** Frequency of markers being affected only by one or two of the three T-cell subsets. Statistical significance was calculated by a paired Student t test. Black circles: staining on fresh PBMCs, Red circles: staining on 6-month frozen PBMCs. The value of each sample is displayed in the figures. ns or unlabeled, not significant, *P<=0.05, **P<=0.01 and ***P<=0.001.

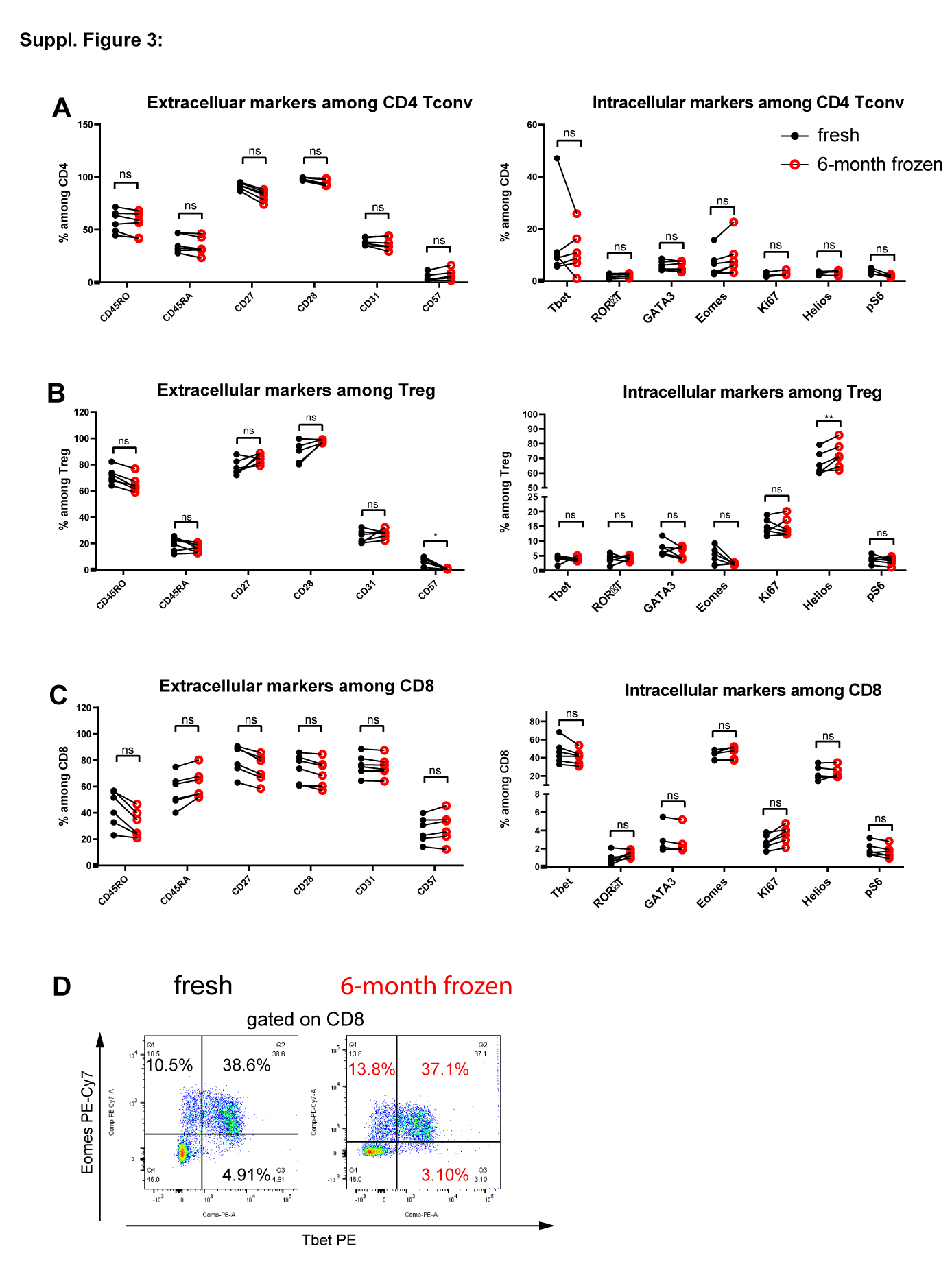

##### Extended Data Figure 3. Six-month cryopreservation does not impair the detection of many extracellular and all tested intracellular T-cell markers.

**A.** Frequency of different unaffected extracellular and intracellular markers among CD4 Tconv in fresh vs. 6-month frozen PBMCs. **B** Frequency of different extracellular and intracellular markers among CD4 Treg in fresh vs. 6-month frozen PBMCs. **C.** Frequency of different unaffected extracellular and intracellular markers among CD8 Tcells in fresh vs. 6-month frozen PBMCs. **D.** Representative dot plots showing the co-expression of intracellular markers Eomes (unaffected) vs. Tbet (unaffected) in fresh (black) or 6-month frozen (red) PBMCs among CD8 T cells. Statistical significance was calculated by a paired Student t test. Black circles: staining on fresh PBMCs, Red circles: staining on 6-month frozen PBMCs. The value of each sample is displayed in the figures. ns or unlabeled, not significant, *P<=0.05, **P<=0.01 and ***P<=0.001.

### Supplementary Table

##### Supplementary Table 1. Different antibodies used at different time points or conditions.

| **Target** | **Fluorochrome** | **Dilution** | **Company** | **Reference** | **Clone** | **Timepoint** |
| --- | --- | --- | --- | --- | --- | --- |
| Fc Blocking Abs | / | 1:50 | BD | 564765 | / | All |
| CD4 | BUV395 | 1:100 | BD | 563550 | SK3 | All |
| CD8 | BUV496 | 1:100 | BD | 564804 | RPA-T8 | All |
| CD25 | PerCP-Cy5.5 | 1:50 | BD | 560503 | M-A251 | Fresh and 9-weeks |
| CD25 | PerCP-Cy5.5 | 1:50 | BioLegend | 302625 | BC96 | 6 months |
| CD27 | APC | 1:50 | BD | 561786 | M-T271 | All |
| CD28 | BUV785 | 1:50 | BioLegend | 302950 | CD28.2 | All |
| CD31 | BV605 | 1:50 | BD | 562855 | WM59 | All |
| CD39 | BV711 | 1:50 | BioLegend | 328228 | A1 | All |
| CD45RA | Pacific Blue | 1:50 | BioLegend | 304123 | HI100 |  |
| CD45RO | PE-CF594 | 1:50 | BD | 562299 | UCHL1 | All |
| CD57 | FITC | 1:50 | BD | 555619 | NK-1 | All |
| CD122 | PE | 1:50 | BioLegend | 339006 | TU27 | All |
| CD127 (IL7R) | V450 | 1:50 | BD | 560823 | HIL-7R-M21 | Fresh and 9-weeks |
| CD127 (IL7R) | BV711 | 1:50 | BioLegend | 351328 | A019D5 | 6 months |
| CD152 (CTLA4) | PE-Cy5 | 1:20 | BD | 561717 | BNI3 | Fresh and 9-weeks |
| CD152 (CTLA4) | PE-Cy7 | 1:20 | BioLegend | 349914 | L3D10 | 6 months |
| CD197 (CCR7) | Pacific Blue | 1:50 | BioLegend | 353210 | G043H7 | Fresh and 9-weeks |
| CD197 (CCR7) | V450 | 1:50 | BD | 560864 | 150503 | 6 months |
| CD278 (ICOS) | BV605 | 1:50 | BioLegend | 313538 | C398.4A | Fresh and 9-weeks |
| CD278 (ICOS) | BV605 | 1:50 | BD | 745100 | DX29 | 6 months |
| CD279 (PD-1) | BV605 | 1:50 | BioLegend | 329924 | EH12.2H7 | Fresh and 9-weeks |
| CD279 (PD-1) | BV605 | 1:50 | BioLegend | 367425 | NAT105 | 6 months |
| KLRG1 | PE-Cy7 | 1:50 | BioLegend | 368614 | 14C2A07 | Fresh and 9-weeks |
| KLRG1 | PE-Cy7 | 1:50 | BioLegend | 138416 | 2F1/KLRG1 | 6 months |
| **Intracellular markers** | | | | | |  |
| FOXP3 | APC | 1:20 | BioLegend | 320114 | 206D | All |
| Phospho S6 | AF488 | 1:20 | CST | 4803S | D57.2.2E | All |
| Helios | Pacific Blue | 1:20 | BioLegend | 137220 | 22F6 | All |
| Ki-67 | FITC | 1:20 | BD | 561165 | B56 | All |
| GATA3 | PE-Cy7 | 1:20 | BD | 560405 | L50-823 | All |
| RORgT | BV650 | 1:20 | BD | 563424 | Q21-559 | All |
| T-bet | PE | 1:20 | BioLegend | 644810 | 4B10 | All |
| Eomes | PE-Cy7 | 1:20 | Thermo Fischer Scientific | 25-4877-42 | WD1928 | All |
| Live/Dead | APC-Cy7 | 1:500 | Thermo Fischer Scientific | L34976 | / | All |

##### Supplementary Table 2. Other flow cytometry reagents used in this work.

| **Reagent** | **Company** | **Reference** |
| --- | --- | --- |
| True-Nuclear Transcription Factor Buffer Set | BioLegend | 424401 |
| Brilliant Stain Buffer | BD | 563794 |
| UltraComp eBeads | eBioscience | 01-2222-42 |
